# *BFA1* has multiple positive roles in directing late mitotic events in *S. cerevisiae*

**DOI:** 10.1101/352427

**Authors:** J Whalen, C Sniffen, S Gartland, M Vannini, A Seshan

## Abstract

The proper regulation of cell cycle transitions is paramount to the maintenance of cellular genome integrity. In budding yeast, the mitotic exit network (MEN) is a Ras-like signaling cascade that effects the transition from M phase to G1 during the cell division cycle in budding yeast. MEN activation is tightly regulated. It occurs during anaphase and is coupled to mitotic spindle position by the spindle position checkpoint (SPoC). Bfa1 is a key component of the SPoC and functions as part of a two-component GAP complex along with Bub2. The GAP activity of Bfa1-Bub2 keeps the MEN GTPase Tem1 inactive in cells with mispositioned spindles, thereby preventing inappropriate mitotic exit and preserving genome integrity. Interestingly, a GAP-independent role for Bfa1 in mitotic exit regulation has been previously identified. However the nature of this Bub2-independent role and its biological significance are not understood. Here we show that Bfa1 also activates the MEN by promoting the localization of Tem1 primarily to the daughter spindle pole body (dSPB). We demonstrate that the overexpression of *BFA1* is lethal due to defects in Tem1 localization, which is required for its activity. In addition, our studies demonstrate a Tem1-independent role for Bfa1 in promoting proper cytokinesis. Cells lacking *TEM1*, in which the essential mitotic exit function is bypassed, exhibit cytokinesis defects. These defects are suppressed by the overexpression of *BFA1*. We conclude that Bfa1 functions to both inhibit and activate late mitotic events.

## INTRODUCTION

In *S. cerevisiae*, the inactivation of the cyclin B homolog Clb2 triggers the transition from mitosis into G1 known as exit from mitosis. This process is regulated by a small-guanine nucleoside triphosphatase (GTPase) signaling cascade known as the mitotic exit network (MEN; Stegmeier and Amon, 2004). Tem1, a small GTPase localized to spindle pole bodies (SPBs), functions at the top of the pathway and activates the downstream Hippo-like kinase Cdc15 (Bardin et al., 2000; Pereira et al., 2000; Asakawa et al., 2001; Lee et al., 2001; Visintin and Amon, 2001). Cdc15 is coordinately recruited to SPBs and activated by both Tem1 and by the Polo protein kinase Cdc5 (Cenamor et al., 1999; Bardin et al., 2000; Pereira et al., 2000; Xu et al., 2000; Molk et al., 2004; Rock and Amon, 2011). Cdc15 in turn activates the Ndr-family kinase Dbf2 (Mah et al., 2001; Visintin and Amon, 2001). The activation of Dbf2 requires an associated factor, Mob1, and is preceded by the recruitment of Mob1 to the MEN scaffold Nud1 at SPBs (Komarnitsky et al., 1998; Luca and Winey, 1998; Mah et al., 2001; Rock et al., 2013). The activation of Mob1-Dbf2 at the SPB and subsequent dissociation of this complex from SPBs results in the activation of the Cdc14 phosphatase, the ultimate effector of the pathway. Cdc14 is held inactive in the nucleolus by its inhibitor Cfi1/Net1. The Mob1-Dbf2 complex causes the release of Cdc14 from Cfi1 during late anaphase, which allows Cdc14 to spread throughout the nucleus and cytoplasm and reach its targets. Activated Cdc14 triggers mitotic CDK inactivation and hence exit from mitosis (Jaspersen et al., 1999; Shou et al., 1999; Visintin et al., 1999).

The regulation of Tem1 activity is important for the integrity of two cell cycle checkpoints: the Spindle Assembly Checkpoint (SAC) and the Spindle Position Checkpoint (SPoC). The SAC acts in metaphase to monitor mitotic spindle – chromosomal attachments before anaphase onset, thereby preventing chromosome missegregation and aneuploidy (reviewed in Musacchio and Salmon, 2007). The SPoC acts in anaphase to monitor alignment of the segregating chromosomes along the mother-bud axis (reviewed in Caydasi et al., 2010). A key role of the two component GTPase activating (GAP) complex Bub2-Bfa1 is to restrain Tem1 activation in the presence of unattached chromosomes or in the case of mitotic spindle misalignment (Alexandru et al., 1999; Fesquet et al., 1999; Fraschini et al., 1999). This, in turn, prevents unscheduled mitotic exit. Indeed, increased levels and activation of Tem1 lead to an increased proportion of cells that bypass the SAC (Chan and Amon, 2009). Therefore it is important to have a thorough understanding of how Tem1 activation is controlled.

The recruitment of Tem1 to SPBs is crucial for its activity. For example, a strain in which Tem1 is mis-localized to the plasma membrane is unable to exit from mitosis (Valerio-Santiago and Monje-Casas, 2011). Conversely, a fusion between Tem1 and the SPB outer plaque component Cnm67 (Tem1-Cnm67) localizes constitutively to both SPBs and leads to bypass of the SPoC (Valerio-Santiago and Monje-Casas, 2011). Thus, Tem1 localization is intertwined with its activation, but how is this process regulated?

Like Tem1, both Bub2 and Bfa1 also localize to SPBs. Bub2 localization is dependent upon Bfa1 localization and Bfa1 is asymmetrically localized to the dSPB from early anaphase through cytokinesis (Pereira et al., 2000). There is mounting evidence that Tem1 and Bfa1 localization to SPBs is interdependent. For example, in cells lacking *BFA1*, Tem1 localizes to a much lesser extent to both SPBs from metaphase to late anaphase, but is clearly present on both SPBs in telophase. However, this does not impair the ability of cells to exit from mitosis in a timely manner (Pereira et al., 2000; Valerio-Santiago and Monje-Casas, 2011). Tem1 localization influences the residence of Bfa1 on the SPBs, although Bfa1 can localize to the dSPB in the absence of Tem1 function (Pereira et al., 2000). In the presence of a Tem1-Cnm67 fusion, Bfa1 is found symmetrically on both SPBs in metaphase and anaphase cells (Valerio-Santiago and Monje-Casas, 2011). Lastly, tethering Bfa1 to the SPBs constitutively using a *SPC72-BFA1* fusion leads to partial bypass of the SPoC and suppresses the localization-defective *tem1-3* allele (Scarfone et al., 2015). These results provide evidence that Bfa1 may have a positive effect on Tem1 localization and therefore, on MEN activation. However, such a role has not been clearly explored thus far.

A MEN-promoting function for Bfa1 would need to be a GAP-independent function. Interestingly, a GAP-independent role for Bfa1 in mitotic exit has been previously described. Bub2 and Bfa1 together increase the intrinsic GAP-activity of Tem1 *in vitro.* Paradoxically, Bfa1 alone appears to inhibit both GTP dissociation and GTP hydrolysis of Tem1 *in vitro*, while having no affect on GDP dissociation (Geymonat et al., 2002). These *in vitro* data would suggest that Bfa1 positively regulates Tem1, once it becomes GTP-bound, and predict that Bfa1, when overexpressed, could activate the MEN. However, the overexpression of *BFA1* instead produces a cell cycle block in anaphase. Interestingly, Ro et al. showed that this terminal arrest was independent of *BUB2*, suggesting that a GAP-independent function of *BFA1* causes the arrest (Ro et al., 2002). Taken together, these data imply that Bfa1 primarily has a negative role in the regulation of Tem1 *in vivo*. However, the BUB2-independent function of *BFA1* on MEN regulation is unknown.

In this study we demonstrate that *BFA1* overexpression using a *GAL-BFA1* allele leads to a defect in Cdc14 activation. We find that this defect stems from an inability of Tem1 to localize correctly to SPBs in the presence of *GAL-BFA1.* Further, we show that the *GAL-BFA1* mitotic exit defect is completely suppressed by co-overexpression of *TEM1.* We confirm that the overexpression of *BFA1* does not affect Mob1-Dbf2 activation during mitotic exit. Interestingly, our data also suggest that Bfa1 may have a MEN-dependent positive function during cytokinesis. These data underscore the positive role for Bfa1 on Tem1 localization and activation, and suggest a novel role for Bfa1 in promoting efficient cytokinesis.

## RESULTS

### The overexpression of *BFA1* causes a defect in Cdc14 activation

Previous studies characterizing the effects of the *GAL-BFA1* allele on mitotic exit demonstrated that these cells delay in anaphase upon galactose induction.

Furthermore, it was known that this severe anaphase delay was not dependent on *BUB2* (Li, 1999; Ro et al., 2002; Figure 1). In order to elucidate the nature of the *BUB2-*independent effects of *BFA1* on anaphase progression, we first sought to determine whether overexpression of *BFA1* affects the release of Cdc14 from its inhibitor Cfi1. In wild type cells undergoing a synchronous cell cycle, the timing of anaphase correlates with the full (nuclear and cytoplasmic) release of Cdc14 from the nucleolus, where it is held inactive by its inhibitor Cfi1. This event is a marker for MEN activation (Jaspersen et al., 1999; Shou et al., 1999; Visintin et al., 1999). We arrested cells in G1 using alpha-factor pheromone and released them into a synchronous cell cycle. We monitored the appearance and disappearance of metaphase and anaphase spindles, as well as Cdc14 localization. Specifically, we analyzed the localization of Cdc14 in anaphase cells. We found that synchronized *GAL-BFA1* cells exhibit a severe anaphase delay and also fail to exhibit full Cdc14 activation and release to the nucleus and cytoplasm (Figure 2A).

**Figure 1.**
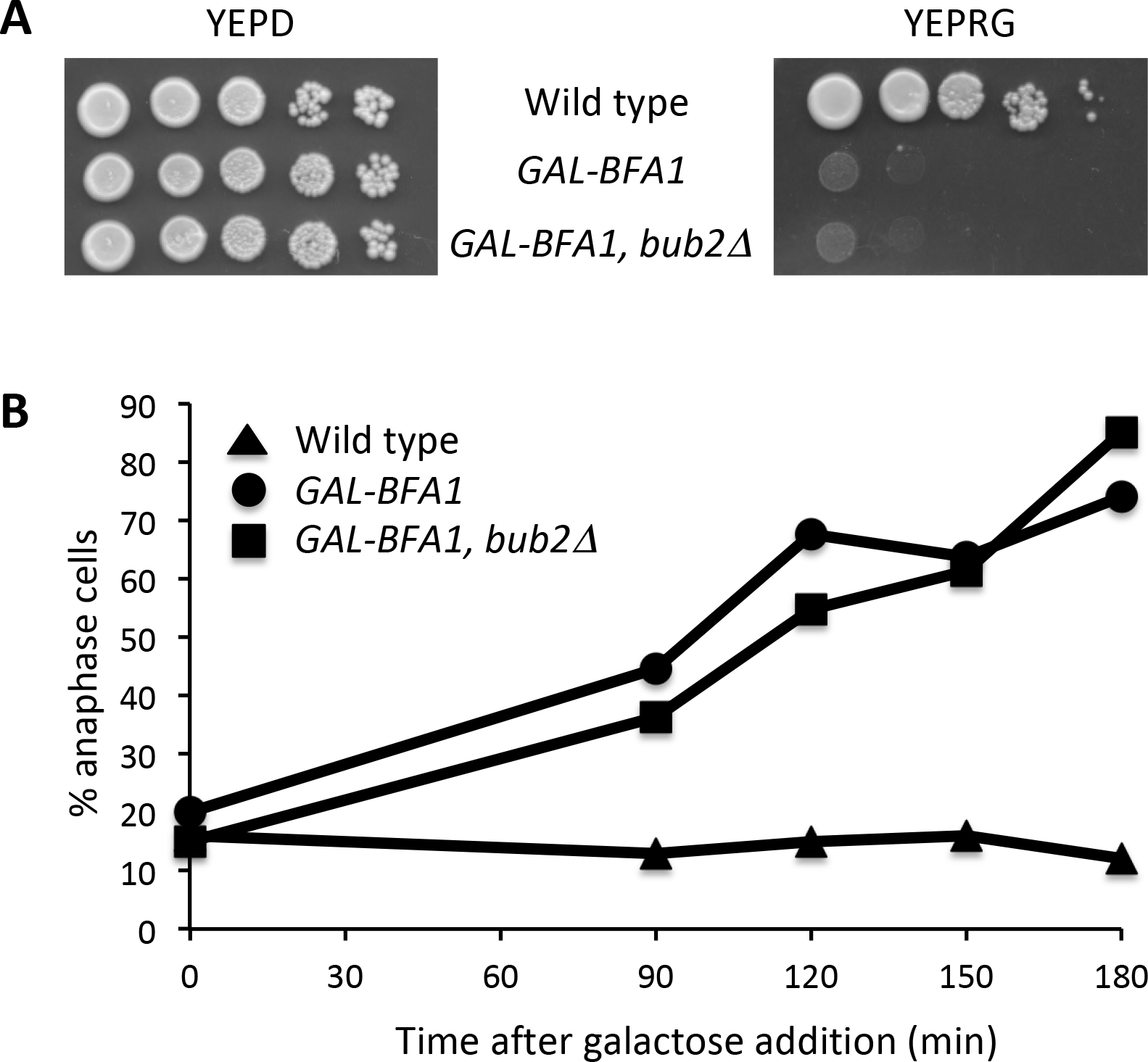
*BFA1* has a *BUB2*-independent late anaphase function. A) Ten-fold serial dilutions of wild type (AS3), *GAL-GFP-BFA1* (AS5), and *GAL-GFP-BFA1 bub2Δ* (AS206) cells were spotted onto YEPD or YEP+Raffinose+Galactose (YEPRG) plates and incubated at 30°C for two days before imaging. B) Galactose was added at time 0 to the log phase, asynchronous, YEP+Raffinose (YEPR) cultures of the indicated genotypes grown at 21°C in order to induce the overexpression of *GAL-GFP-BFA1*. Samples were taken at the indicated times and processed for tubulin immunofluorescence. The percentage of anaphase cells was determined at each time point (n = 100 – 200 cells).

**Figure 2.**
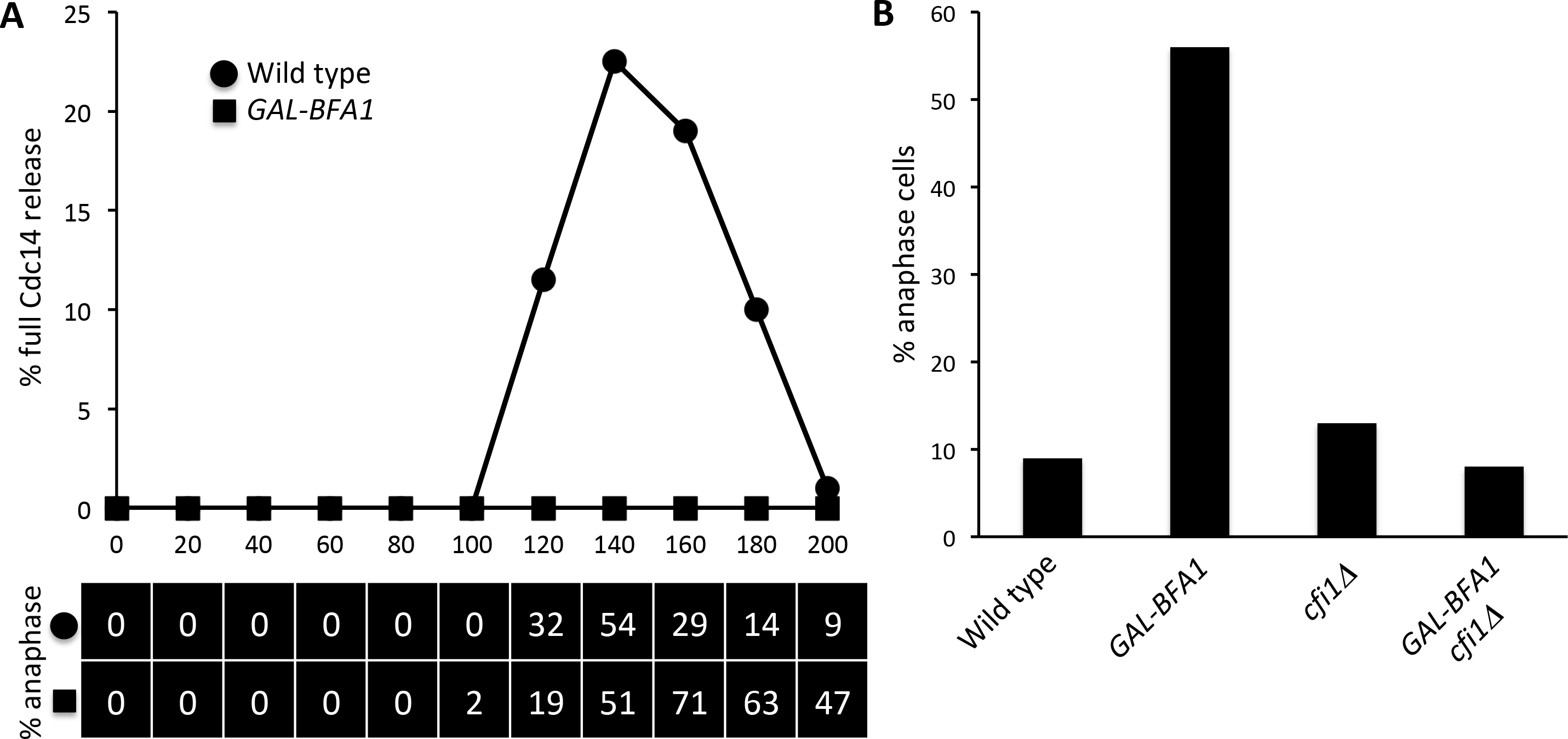
Cdc14 activation is impaired in *GAL-BFA1* cells. A) Log phase cultures of wild type (AS138) and *GAL-BFA1* (AS24) cells carrying a *CDC14-3HA* fusion growing at 21°C in YEPR were arrested in G1 using alpha-factor pheromone (5 μg /mL). At 1.5 hours into the arrest, galactose (GAL) was added to induce expression of *GAL-BFA1.* Cells were released from the G1 arrest after 3 hours into YEPRG. Cells were collected at the times indicated to process samples for tubulin and Cdc14-HA immunofluorescence. The percentage of cells with anaphase spindle morphology was quantified for each time point and is indicated in the chart below the graph. The percentage of cells with Cdc14 localized to the nucleus and cytoplasm (full Cdc14 release) was quantified in wild type (circles) and *GAL-BFA1* (squares) cells (N = 100 - 200 cells). B) Log phase wild type (AS3), *GAL-BFA1* (AS5), *cfilΔ* (AS121), and *cfilΔ GAL-BFA1* (AS141) were grown in YEPRG at 21°C for three hours to induce the overexpression of *BFA1* and samples were collected and processed for tubulin immunofluorescence. The percentage of cells with anaphase spindle morphology was quantified (N = 200 cells).

We next examined whether the deletion of *CFI1* would ameliorate the anaphase delay of *GAL-BFA1* cells. While the induction of *BFA1* overexpression in *GAL-BFA1* cells led to a significant anaphase delay, *GAL-BFA1 cfi1Δ* cells resembled wild type cells and did not have anaphase defects (Figure 2B). Taken together, these results demonstrate that Cdc14 activation is blocked by *GAL-BFA1* and that the effects of *BFA1* overexpression can be attenuated by Cdc14 hyperactivation.

### *BFA1* overexpression causes de-localization of Tem1 from SPBs

We were interested in determining the specific step in the MEN that was perturbed by the *GAL-BFA1* allele. There is prior evidence that Tem1 and Bfa1 localization to SPBs is at least partially interdependent (Pereira et al., 2000; Valerio-Santiago and Monje-Casas, 2011; Scarfone et al., 2015). Therefore, we hypothesized that both Tem1 and Bfa1 localization were impaired when *BFA1* was overexpressed. We first examined the localization of the *GAL-GFP-BFA1* allele. We examined anaphase cells and were surprised to find that in the presence of galactose, Bfa1-GFP is concentrated at the dSPB, as it is in wild type cells (Pereira et al., 2000). However, we also noted that significant cytoplasmic signal is present in these cells (Figure 3A).

**Figure 3.**
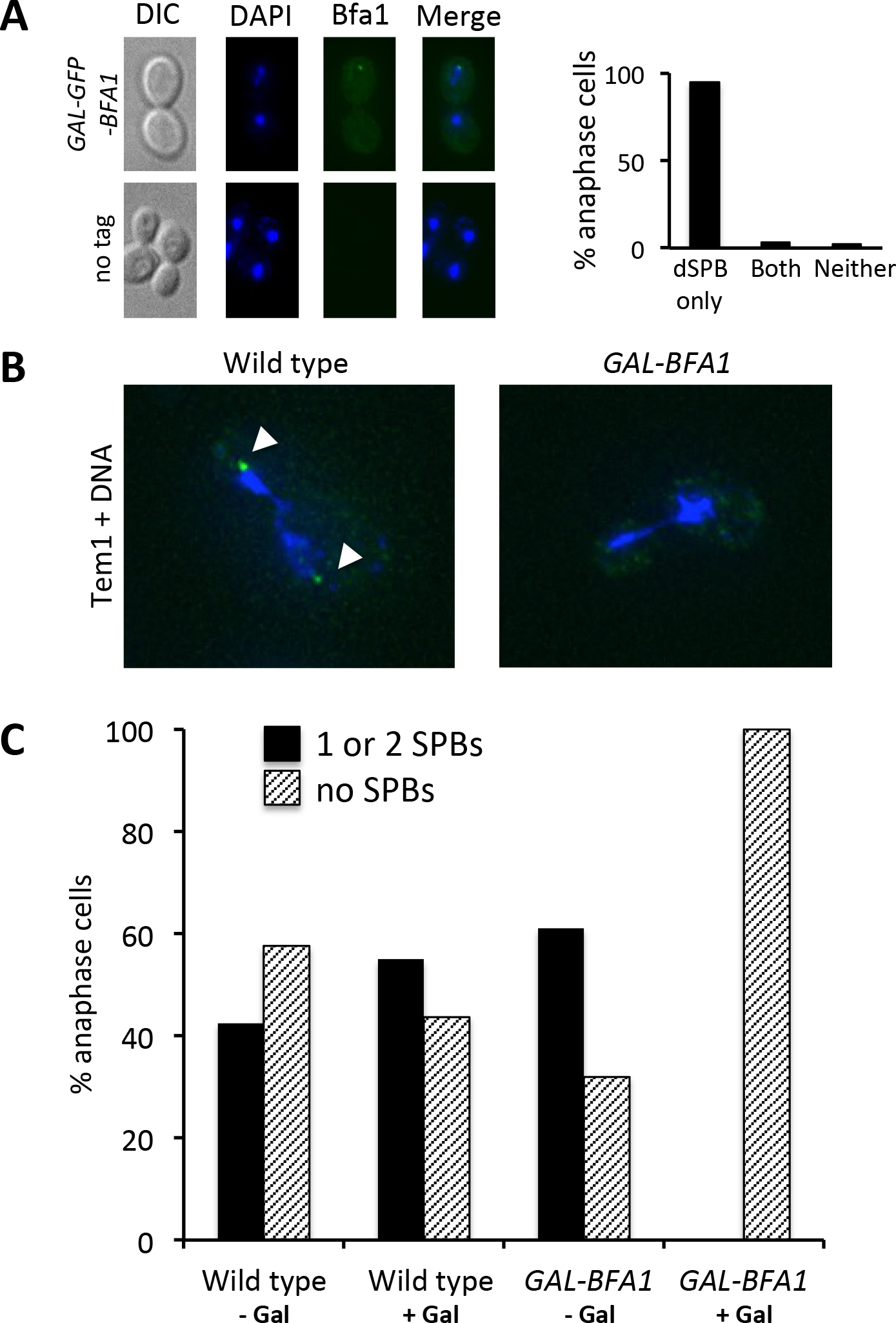
Tem1 localization to SPBs is perturbed in *GAL-BFA1* cells. A) A *GAL-GFP-BFA1* strain (AS5) was grown to log phase in YEPR at 21°C and the cells were arrested in G1 with alpha-factor pheromone (5 μg/mL). Cells were released from the arrest in YEPRG and the overexpression of *BFA1* was induced for a total of 4.5 hours. Cells were fixed with paraformaldehyde prior to imaging. Representative anaphase cells are shown. The percentage of anaphase cells with Bfa1 localized to the dSPB, both the dSPB and the mSPB (both), or to neither SPB was quantified (n = 133). Asynchronous no tag wild type (AS3) cells were also imaged. B and C) Wild type (AS15) and *GAL-BFA1* strains (AS79) containing the Tem1-GFP fusion were grown in YEPR (− Gal) or YEPRG (+ Gal) media for three hours at 21°C before paraformaldehyde fixation and imaging. Anaphase cells were identified by DAPI morphology and 50 to 100 cells were analyzed in each condition. Representative anaphase cells are shown in B). Tem1-GFP is displayed in green and the DNA is shown in blue. Tem1-GFP localization to SPBs is indicated by the white arrowheads. C) The percentage of anaphase cells in each sample with Tem1-GFP localized to one or both SPBs (black bars) or delocalized throughout the cell (hatched bars) was plotted.

Given the significant cytoplasmic localization of Bfa1 in *GAL-BFA1* cells, we reasoned that Tem1 could be dragged off of the SPB and into the cytoplasm in these cells. Indeed we found that Tem1-GFP was delocalized from SPBs in 100% of cells when *BFA1* was overexpressed by galactose addition. In contrast, when these same cells were cultured in the presence of raffinose alone, a majority of the cells had Tem1 localized to one or two SPBs (Figure 3). These results demonstrate that Tem1 localization is defective in *GAL-BFA1* cells.

### Tem1 re-localization to SPBs does not fully suppress *GAL-BFA1* mitotic exit defects

Now that we had determined that Tem1 localization is affected in *GAL-BFA1* cells, we hypothesized that we could suppress this defect by artificially re-localizing Tem1 to SPBs in *BFA1* overexpressing cells. In the presence of a Tem1-Cnm67 fusion, Bfa1 is found symmetrically on both SPBs in metaphase and anaphase cells (Valerio-Santiago and Monje-Casas, 2011). Therefore we analyzed the kinetics of Cdc14 release from Cfi1 in the nucleolus and the kinetics of mitotic exit in *GAL-BFA1 TEM1-eGFP-CNM67* cells. In this experiment, we analyzed the localization of Cdc14 in anaphase cells to the nucleolus, the nucleus, and the nucleus and cytoplasm. This allowed us to distinguish between cells with inactive Cdc14 (sequestered in the nucleolus), Cdc14 released by the Cdc Fourteen Early Anaphase Release (FEAR) network (nucleus only; reviewed in Rock and Amon 2009), and Cdc14 released in late anaphase by the MEN into the nucleus and the cytoplasm. We found that the partial release of Cdc14 to the nucleus that occurs in early anaphase due to FEAR pathway activity was unaffected in *GAL-BFA1* cells. However, as shown previously these cells accumulated in anaphase and 63% of anaphase cells displayed Cdc14 sequestration in the nucleolus, indicating that the MEN was inhibited in these cells (Figure 4F). We found that cells containing both *GAL-BFA1* and *TEM1-eGFP-CNM67* were able to exit from mitosis in a slightly timelier manner than *GAL-BFA1* cells, though they were by no means able to progress like wild type cells (Figure 4]). However, we observed only a partial suppression of the anaphase delay, as 31% of anaphase cells in the *GAL-BFA1 TEM1-eGFP-CNM67* background still had Cdc14 sequestered in the nucleolus (Figure 4F). Importantly this partial suppression was not due to de-localization of Tem1-Cnm67 in the *GAL-BFA1* background, as the Tem1-eGFP-Cnm67 protein was observed at the dSPB or at both mother and daughter SPBs regardless of *GAL-BFA1* induction. Interestingly, we observed that in the presence of the *GAL-BFA1* allele, a larger fraction of anaphase cells exhibited a bias for Tem1-Cnm67 dSPB localization than in *TEM1-CNM67* cells that did not overexpress *BFA1*, which further suggests that Bfa1 influences Tem1 localization to SPBs (Figure 4G-H).

**Figure 4.**
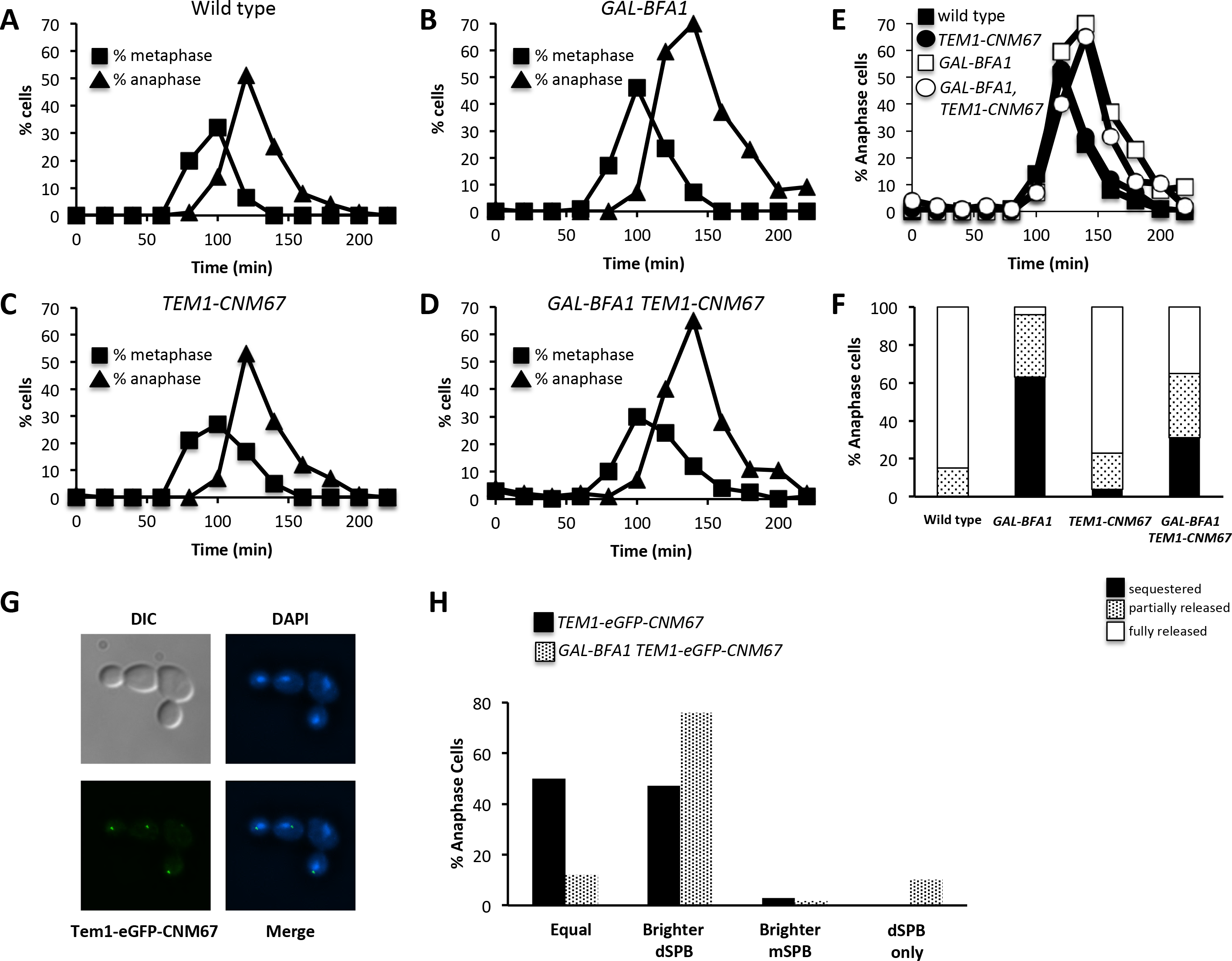
Restoration of Tem1 localization is not sufficient to suppress the effects of *GAL-BFA1*. Log phase YEPR cultures of wild type (AS138), *GAL-BFA1* (AS24), *TEM1-eGFP-CNM67* (AS109), and *GAL-BFA1 TEM1-eGFP-CNM67* (AS188) cells carrying a CDC14-3HA fusion were arrested with alpha-factor pheromone (5 μg/mL) at 21°C. At 1.5 hours into the arrest, galactose was added. The cells were released from the arrest after 3 hours into YEPRG media. Cells were collected at the times indicated to process for tubulin and CDC14-HA immunofluorescence. A-D) The percentage of cells with metaphase (squares) and anaphase (triangles) spindle morphology was quantified for each time point (N= 100-200 cells). E) The percentage of anaphase cells at each time point from (A) - (D) was plotted for comparison. F) The percentage of anaphase cells of the genotypes indicated with Cdc14 sequestered (black), partially released (dotted) or fully released (white) was plotted (N= 100 cells). G) *TEM1-eGFP-CNM67* (AS19) and *GAL-BFA1 TEM1-eGFP-CNM67* (AS36J cells were grown at 21°C in YEPRG for three hours and cells were imaged after paraformaldehyde fixation. Representative *GAL-BFA1 TEM1-eGFP-CNM67* cells are shown. H) The percentage of *TEM1-eGFP-CNM67* (black bars) or *GAL-BFA1 TEM1-eGFP-CNM67* (stippled bars) anaphase cells with Tem1-eGFP-CNM67 localized equally on both SPBs, brighter on the dSPB, brighter on the mSPB, or located only on the dSPB was quantified (N = 100 cells).

### *GAL-TEM1*, but not *TEM1-2μ*, can fully suppress *GAL-BFA1* mitotic exit defects

We considered two hypotheses for the partial suppression of *GAL-BFA1* by *TEM1-CNM67*. One possibility was that the *TEM1-CNM67* allele provides low levels of activated Tem1 and higher levels are needed to fully bypass *GAL-BFA1*. A second, non-mutually exclusive possibility was that *GAL-BFA1* has TEM1-independent effects on MEN activation. We decided to test this first possibility by examining the suppression of *GAL-BFA1* by two different alleles of *TEM1: TEM1-2μ and GAL-TEM1.* We predicted that *TEM1-2μ* would also exhibit a partial bypass of *GAL-BFA1* because this allele also has lower levels of activated Tem1. Conversely, we expected to observe complete suppression of the effects of *GAL-BFA1* by the *GAL-TEM1* allele because this allele exhibits high levels of activated Tem1 (Chan and Amon, 2009). We synchronized cells and utilized spindle and nuclear morphology were used to assess the timing of metaphase, anaphase and mitotic exit. We found that the anaphase delay phenotype observed in *GAL-BFA1* was not suppressed in *GAL-BFA1 TEM1-2μ* cells, which contained the *TEM1* gene on a multicopy plasmid (Figure 5). In contrast, the timing of mitotic exit in *GAL-BFA1 GAL-TEM1* cells overexpressing both *BFA1* and *TEM1* was indistinguishable from wild type cells (Figure 6). We noted that the extent of the anaphase delay observed in *GAL-BFA1* cells varied depending upon the type of media utilized and the temperature at which cells were cultured (compare 4B, 5B, and 6B). However regardless of culturing conditions, these data support the hypothesis that defects in Tem1 activation are the primary reason that *GAL-BFA1* cells cannot activate the MEN.

**Figure 5.**
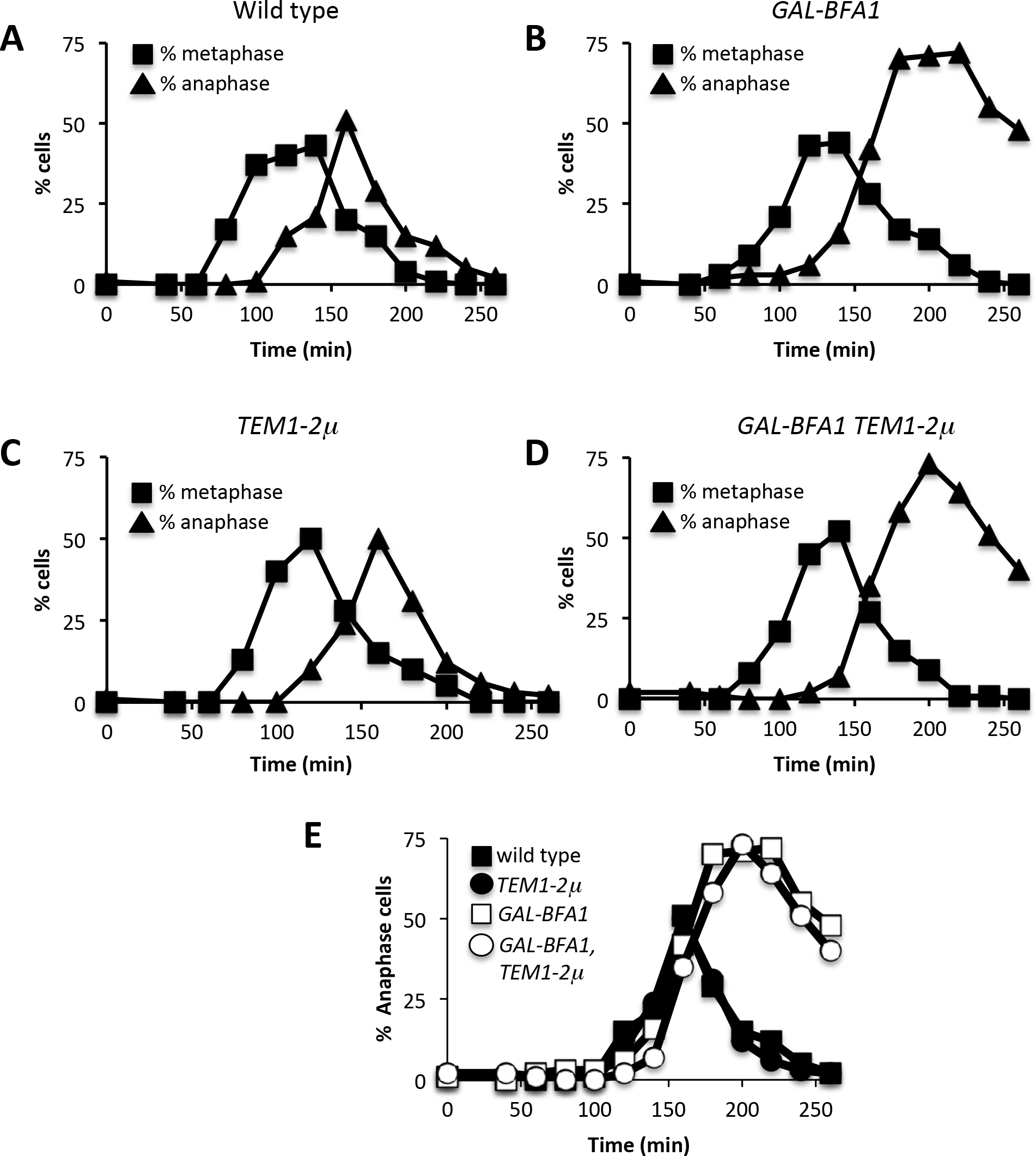
The *TEM1-* 2μ allele does not suppress *GAL-BFA1.* A – D) Log phase cultures of wild type + YEP 2μ (empty vector; AS207), wild type + *TEM1*-2μ (AS209), *GAL-BFA1* + YEP 2μ (AS211), and *GAL-BFA1* + *TEM1-* 2μ (AS213) were arrested using alpha-factor pheromone (5 μg/mL) at 21°C in SC-Leu + Raffinose media. At 1.5 hours into the arrest, galactose was added to the cells to induce the expression of *GAL-BFA1.* Cells were released from the arrest into SC-Leu + Raffinose +Galactose media after 3 hours. Samples of cells were collected at the indicated times and processed for tubulin immunofluorescence. The percentages of metaphase and anaphase cells were determined at each time point (N=100-200). E) The percentage of anaphase cells at each time point from (A) – (D) was plotted for comparison.

**Figure 6.**
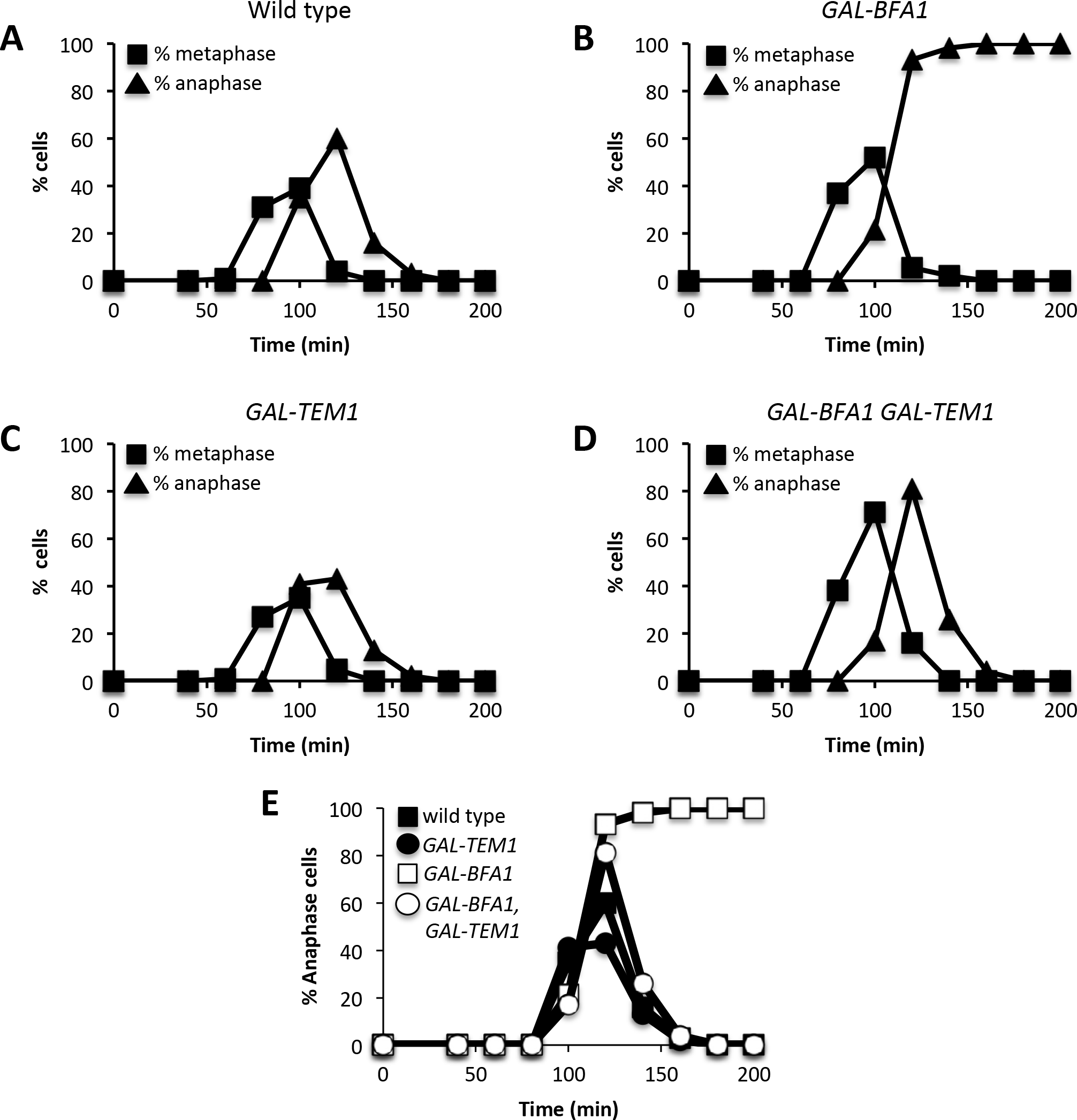
The *GAL-TEM1* allele displays robust suppression of *GAL-BFA1.* A – D) Log phase wild type (AS3), *GAL-BFA1* (AS413), *GAL-TEM1* (AS401), and *GAL-BFA1 GAL-TEM1* (AS414) cells were arrested in YEPR media at 25°C using alpha-factor pheromone (5 μg/mL). At 1.5 hours into the arrest, galactose was added to the cells to induce the expression of *GAL-BFA1* and/or *GAL-TEM1.* Cells were released from the arrest into YEPRG media after 3 hours. Samples of cells were collected at the indicated times and were processed for tubulin immunofluorescence. The percentages of metaphase and anaphase cells were determined at each time point (N=100-200). E) The percentage of anaphase cells at each time point from (A) – (D) was plotted for comparison.

### Dbf2 localization to the budneck is enhanced by *GAL-BFA1*

Our prior data strongly suggested that the mislocalization and faulty activation of Tem1 in *GAL-BFA1* cells resulted in a failure to activate the MEN. However, in order to confirm that the *GAL-BFA1* allele did not impart any *TEM1-* independent mitotic exit defects, we sought to analyze MEN activation in *GAL-BFA1* cells that lacked Tem1 activity. The deletion of *TEM1* confers a lethal phenotype. In order to keep *tem1Δ* cells alive, the hyperactive *CDC15-UP* allele, in which *CDC15* is overexpressed using a GAL-inducible promoter, was utilized (Rock and Amon, 2011). We examined the localization of Dbf2-eGFP to SPBs as a proxy for MEN activation in *tem1Δ CDC15-UP* cells. Dbf2 is known to localize to SPBs during late anaphase following Cdc15 activation, but the protein also localizes to the budneck where it activates the process of cytokinesis (Stegmeier and Amon, 2004; Frenz et al., 2000; Meitinger et al., 2010; Oh et al., 2012; Meitinger et al., 2013). In order to fully assess the effects of *GAL-BFA1* on Dbf2 localization, we quantified the number of anaphase cells with Dbf2-eGFP at the SPBs, at the budneck, at both of these structures, and at neither of these structures (Figure 7A).
In large budded wild type cells containing one Spc42-mCherry SPB dot distributed in mother and daughter cells, Dbf2-eGFP is found at the SPBs in 42.8% of cells (SPB only + SPB and budneck; Figure 7A). In *tem1Δ CDC15-UP* and *tem1Δ CDC15-UP GAL-BFA1* cells Dbf2-eGFP is found at SPBs in 57.6% and 73.3% of cells respectively (SPB only + SPB and Budneck; Figure 7A). These data indicate that the overexpression of *BFA1* does not have TEM1-independent mitotic exit effects.

**Figure 7.**
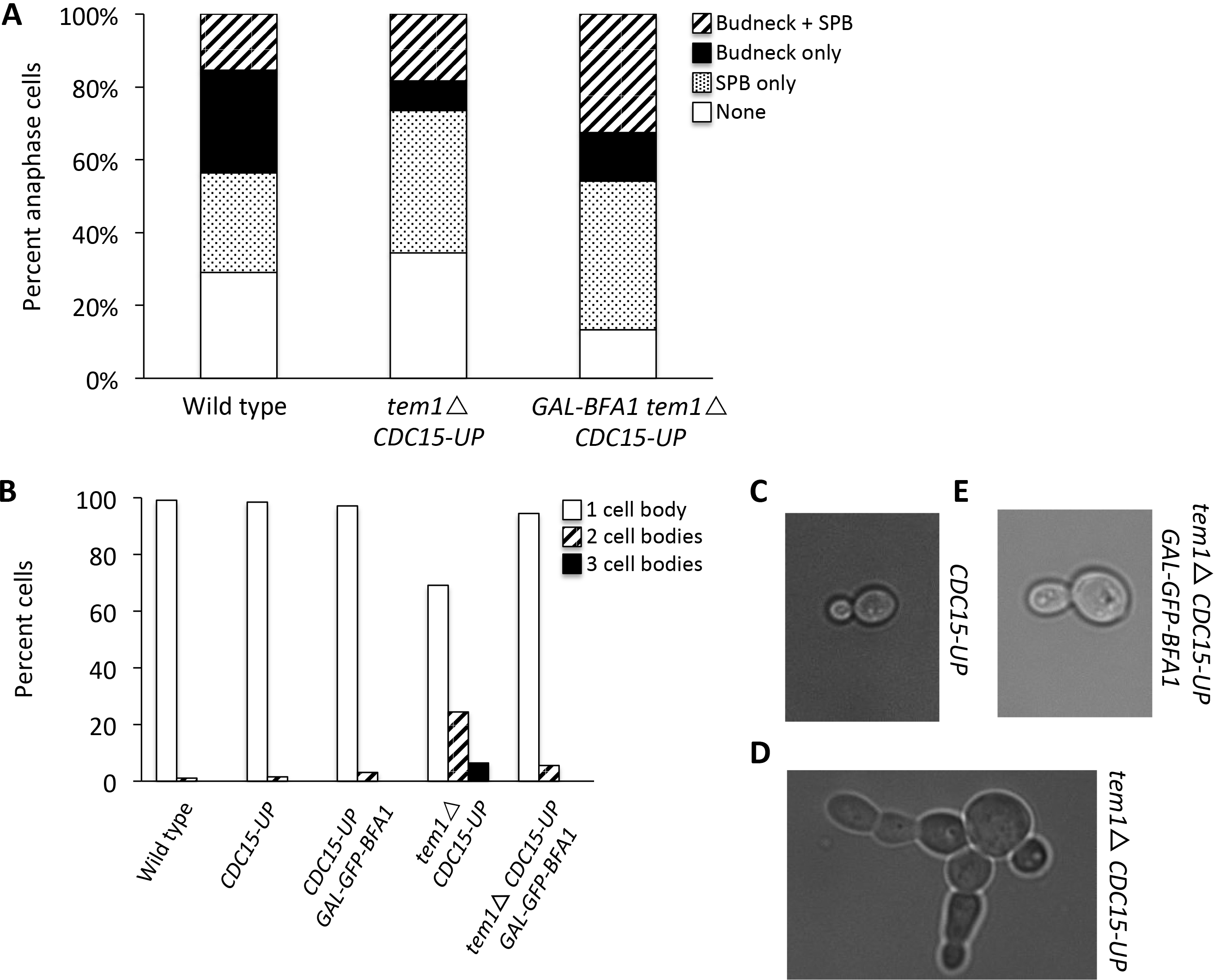
Dbf2 budneck localization and cytokinesis defects in *temlΔ* cells are ameliorated by *GAL-BFA1.* Log phase samples of wild type (AS358), *teml Δ CDC15-UP* (AS431), and *teml Δ CDC15-UP GAL-BFA1* (AS394) strains harboring Dbf2-eGFP and Spc42-mCherry fusions grown at 23°C in SC+RG media were collected and imaged. A) Large budded cells with Spc42 distributed in mother and daughter cells were identified and the percentage of cells with Dbf2-eGFP localized only to one or more spindle pole body (SPB only), localized only to the budneck (Budneck only), to SPBs and to the budneck (Budneck + SPB), or delocalized (None) was quantified in each strain (N = 120 cells). B) Wild type (AS3); *CDC15-UP* (AS118); *CDC15-UP GAL-GFP-BFA1* (AS119); *teml Δ CDC15-UP* (AS259); and *teml Δ CDC15-UP GAL-GFP-BFA1* (AS493) cells were grown to log phase in SC+RG media at 23°C, fixed in formaldehyde, and subjected to brief sonication. The percentage of cells with one cell body, two connected cell bodies, or three connected cell bodies was quantified (n = 200). Representative DIC microscopic images *CDC15-UP* (C); *tem1Δ CDC15-UP* (D); and *tem1 Δ CDC15-UP GAL-GFP-BFA1* (E) cells are shown.

Unexpectedly, when we examined the budneck localization of Dbf2-eGFP in these same cells, we found that Dbf2-eGFP budneck localization is impaired in *tem1Δ CDC15-UP* anaphase cells. Specifically, wild type cells exhibit budneck localization in 43.7% of cells, while *tem1Δ CDC15-UP* cells only have Dbf2-eGFP at the budneck in 26.4% of cells (Budneck only + SPB and Budneck; Figure 7A). In *tem1Δ CDC15-UP GAL-BFA1* cells, however, the fraction of anaphase cells with Dbf2-eGFP localized to the budneck is returned to 45.8% (Budneck only + SPB and Budneck; Figure 7A). These data indicated that, despite the fact that *tem1Δ CDC15-UP* cells display normal cell cycle kinetics and Dbf2 kinase activity during anaphase (Rock and Amon, 2011), the cytokinesis-specific role of Dbf2 was disrupted. They also suggested that the overexpression of *BFA1* could suppress these defects in *tem1Δ CDC15-UP* cells.

To further explore whether cytokinesis defects were present in *tem1Δ CDC15-UP* cells, and to determine whether the deletion of *TEM1* or the overexpression of *CDC15* were responsible for these defects, we analyzed the cellular morphology of these cells following a brief sonication. This allows the separation of cell clumps without disrupting cell walls. We found that less than 2% of wild type and *CDC15-UP* cells exhibited cytokinesis defects where two or more cell bodies remained connected after brief sonication (Figure 7B, 7C). In contrast, 31% of *tem1Δ CDC15-UP* cells exhibited chained cell morphology where two or three cell bodies remained connected (Figure 7B, 7D).

Furthermore, we observed that the cytokinesis defects of *tem1Δ CDC15-UP* cells were largely suppressed in the presence of *GAL-BFA1*. In *tem1Δ CDC15-UP GAL-GFP-BFA1* cells, only 5.5% of cells exhibited chains (Figure 7B, 7E). These data imply that increased levels of *BFA1* promote cytokinesis by a Tem1-independent mechanism.

## DISCUSSION

The activation of the MEN is essential for the destruction of mitotic CDK activity at the M phase to G1 transition. Several components of the MEN, as well as the MEN regulators Bfa1 and Bub2, localize to either one or both SPBs (reviewed in Stegmeier and Amon, 2004). Furthermore, mutations or conditions that perturb SPB localization of Tem1, Cdc15, Mob1, and Dbf2 have been shown to affect timely mitotic exit as well as mitotic checkpoint functions (reviewed in Musacchio and Salmon, 2007; reviewed in Scarfone and Piatti, 2015). Therefore, it is important to deepen our understanding of how the localization of these components to SPBs is regulated. Our findings highlight a positive role for Bfa1 in the localization of Tem1, which is required for MEN activation. In addition, we show that Bfa1 promotes efficient cytokinesis by an unknown mechanism. These data provide another example of how the temporal coupling between mitotic exit and cytokinesis is established in budding yeast.

### *BFA1* overexpression inhibits MEN activity but not FEAR activity

The well-established role of Bfa1 is to negatively regulate the MEN by stimulating Teml’s GTPase activity in complex with the enzymatic component Bub2 (Pereira et al., 2000; Geymonat et al., 2003). However, increasing cellular Bfa1 protein levels using a *GAL-BFA1* allele prevents mitotic exit even in the absence of Bub2 (Ro et al., 2002 and this study). We showed that in *GAL-BFA1* cells there is no MEN-induced release of Cdc14 observed and the cells delay in anaphase. The anaphase delay phenotype is suppressed by hyperactivation of Cdc14 using the *cfi1Δ* allele (Shou et al., 1999; Visintin et al., 1999; Figure 2B). These results confirm that there is a Bub2-independent role for Bfa1 in the regulation of the MEN, which functions to promote Cdc14 release from the nucleolus.

The anaphase delay phenotype in *GAL-BFA1* cells was variable depending upon temperature. For example, we observed that in rich media at 25°C, 100% of cells arrested anaphase (Figure 6B). However, at 21°C in rich media, *GAL-BFA1* cells exhibit a 40 minute delay in anaphase with 63% of anaphase cells containing Cdc14 sequestered in the nucleolus, but eventually these cells break down their spindles (Figure 4B). It is not clear why the phenotype is variable, but we speculate that the overexpression of *BFA1* is less severe in cells that are growing more slowly and therefore have a prolonged metaphase. We noted that the FEAR-mediated release of Cdc14 was not affected by *GAL-BFA1* (Figure 4F). It is possible that cell-to-cell differences in FEAR network component levels and activity produced especially high levels of FEAR-dependent Cdc14 activation, which allowed some anaphase cells to exit from mitosis despite high levels of MEN-inhibition by *GAL-BFA1*. A FEAR-dependent mitotic exit despite MEN-inhibition has been previously shown in cells with mispositioned anaphase spindles. Falk *et al.* demonstrated that a small number of otherwise wild type cells exit from mitosis inappropriately, even in the presence of mispositioned spindles. However, this inappropriate mitotic exit is completely prevented in cells where the FEAR network has been inactivated by the deletion of the FEAR components *SPO12* or *SLK19* (Falk *et al.*, 2016). If FEAR activity were to be a prerequisite for a proportion of *GAL-BFA1* anaphase cells growing at 21°C to exit from mitosis, the deletion of *SPO12* in these cells would be predicted to extend the anaphase delay.

### *BFA1* overexpression does not inhibit MEN through symmetric SPB localization

Bfa1, when present in the cell at endogenous levels, localizes preferentially to the dSPB (Pereira et al., 2000). When the protein is overexpressed, we see that a significant proportion of GFP-Bfa1 is present at the dSPB and also in the cytoplasm (Figure 3A). Previous studies have suggested that a shift from symmetric Bfa1-Bub2 SPB localization to asymmetric localization pattern precedes the activation of Tem1 and mitotic exit (Pereira et al., 2000, 2001; Molk et al., 2004; Fraschini et al., 2006; Maekawa et al., 2007; Geymonat et al., 2009; Kim et al., 2012). In addition, Cdc5 inactivates Bfa1 in anaphase by phosphorylating Bfa1 (Hu et al., 2001; Geymonat et al., 2003). Therefore, the overexpression of *BFA1* could potentially overwhelm the Cdc5-dependent inhibitory phosphorylation of the protein, and thereby contribute to Bfa1 hyperactivity. However, our finding that GFP-Bfa1 localized asymmetrically to the dSPB in *GAL-BFA1* cells argues against this, as the Bfa1-4A mutant, in which the four Cdc5 phosphorylation sites in Bfa1 are mutated, displays a symmetric localization pattern (Kim et al., 2012). This observation further suggested that the block in MEN activation in the presence of increased Bfa1 protein was not due to increased Bfa1 inhibitory activity.

### Bfa1 promotes Tem1 SPB localization

Several lines of evidence indicate that *BFA1* overexpression deters mitotic exit by preventing proper localization of Tem1 to the dSPB. First, we show that Tem1 is delocalized from SPBs in *GAL-BFA1* strains (Figure 3B, 3C). Tem1 localization to SPBs is likely to be a pre-requisite for mitotic exit. Tethering the protein to the plasma membrane using a Tem1-CAAX fusion blocks mitotic exit. On the other hand, the fusion of *TEM1* to *CNM67*, which encodes a component of the outer plaque of the SPB, causes mitotic to occur even in cells that have mispositioned anaphase spindles (Valerio-Santiago and Monje-Casas, 2011). We demonstrated that artificially tethering Tem1 to SPBs constitutively using the *TEM1-CNM67* allele leads to a partial suppression of *GAL-BFA1* lethality (Figure 4). This finding confirms that removal of Tem1 from the SPBs is a key mechanism by which the overexpression of *BFA1* blocks mitotic exit and argues for a positive function for Bfa1 in mitotic exit regulation.

Although mitotic exit did proceed to some extent in *GAL-BFA1 TEM1-CNM67* cells, a significant proportion of anaphase cells (31%) still had Cdc14 sequestered in the nucleolus (Figure 4E). We propose that this is due to the weak allele strength of the *TEM1-CNM67* allele. Using a checkpoint bypass assay, Chan and Amon previously determined that hyperactive alleles of *TEM1* confer differing levels of MEN hyperactivation. Specifically, the *GAL-TEM1* allele produced the highest MEN activation level in this assay, while both *TEM1-CNM67* and *TEM1-2micron* alleles produced lower levels of MEN activation (Chan and Amon, 2009). In keeping with this, we observed that only the presence of *GAL-TEM1* completely abolished *GAL-BFA1* inhibition of mitotic exit (Figure 6). In addition, we found that *GAL-BFA1* posed no impact on the SPB localization of the downstream MEN kinase Dbf2 when the effects on Tem1 were bypassed. Specifically, cells lacking *TEM1* and kept alive using a hyperactive *CDC15-UP* allele showed normal recruitment of Dbf2 to SPBs both in the presence and absence of overexpressed *BFA1* (Figure 7A). Together, these results support the conclusion that Tem1 mis-localization is the central mitotic exit defect in *GAL-BFA1* cells.

Why is does the *TEM1-CNM67* allele display lower levels of activity, despite the fact that Tem1 localization to SPBs is a prerequisite for its activation? One reason could be that the Tem1-Cnm67 fusion at the SPBs is subject to increased GAP activity, which prevents the protein from being highly active. While we acknowledge that this is possible, we do not favor this explanation because *TEM1-CNM67 bfa1Δ* cells, in which the GAP is inactivated, display the same SPoC bypass phenotype as *TEM1-CNM67* cells (Valerio-Santiago and Monje-Casas, 2011). These data indicate that GAP inactivation does not enhance the activity of *TEM1-CNM67* allele.

The *TEM1-CNM67* allele displays a significantly more symmetric SPB localization than the wild type *TEM1*, and causes endogenous Bfa1 protein to localize in a symmetric manner (Valerio-Santiago and Monje-Casas, 2011; Figure 4G). Here, we showed that when *BFA1* is overexpressed, Bfa1, which is concentrated in high amounts at the dSPB, led to a largely asymmetric localization of the Tem1-Cnm67 chimeric protein to the dSPB (Figure 4G). This confirms that Tem1-Cnm67 and Bfa1 interact at the SPB even when Bfa1 is present at high levels. This also provides further evidence that Bfa1 acts as a receptor for Tem1 at the dSPB.

### How does overexpressed *BFA1* disrupt Tem1 localization to SPBs?

Tem1 requires Bfa1 for efficient loading onto SPBs, though the protein can be found on SPBs at low levels even in the absence of *BFA1* (Pereira et al. 2000; Valerio-Santiago and Monje-Casas 2011). Although it is not known exactly how Tem1 associates with SPBs, Tem1 itself can associate with both Bfa1 and Nud1 in two-hybrid binding assays (Kim et al., 2008; Valerio-Santiago and Monje-Casas, 2011). Bfa1 itself binds in a complex at the SPB outer plaque with both Nud1 and Spc72 (Gruneberg et al., 2000; Gryaznova et al., 2016). It has been suggested that in the absence of *BFA1*, Tem1 may bind to Nud1 (Valerio-Santiago and Monje-Casas, 2011). Therefore, it is possible that the large amounts of Bfa1 that are present saturate the outer plaque, and specifically Nud1, and prevent the localization of Tem1 to this organelle. However, we favor the idea that the strong association between cytoplasmic Bfa1 in the *GAL-BFA1* allele and Tem1 titrates Tem1 away from the SPB. Notably, in *tem1Δ CDC15-UP GAL-BFA1* cells, Dbf2-eGFP is observed at SPBs, presumably bound to its receptor phospho-Nud1 (Rock et al., 2013; Figure 7). This indicates that increased levels of Bfa1 on the dSPB do not prevent the association of other MEN components with the Nud1 scaffold on the outer plaque.

### Bfa1 serves as both an activator and an inhibitor of late M phase events

The essential mitotic exit function of Tem1 is to recruit Cdc15 to the SPB by an unknown mechanism (Rock et al., 2011). Previous work has demonstrated that the fusion of *BFA1* with *TEM1* or with the SPB outer plaque component *SPC72* leads to increased, rather than decreased, MEN activation by increasing the residence time of Tem1 at SPBs (Valerio-Santiago and Monje-Casas, 2011; Scarfone et al., 2015). Our results are consistent with the notion that Bfa1 can enhance Tem1 activity and MEN activation by bringing Tem1 to the SPB, and that preventing Tem1 localization by toggling the cellular levels of Bfa1 can conversely inhibit MEN.

The Septation Initiation Network (SIN) in *S. pombe* is homologous to the MEN, but controls septation and cytokinesis rather than mitotic cyclin inactivation (reviewed in Krapp and Simanis, 2008). The Tem1 homolog Spg1p and the Bfa1 homolog Byr4p also localize to the SPB in an asymmetric manner (Cerutti and Simanis, 1999; Li et al., 2000). Byr4p serves as a bridge on the SPB between Spg1p and the Bub2 homolog Cdc16p, which is the enzymatic component of the Byr4p-Cdc16p two-component GAP complex (Furge et al., 1998). It was previously shown that overexpression of Byr4p blocks SIN activation and cytokinesis due to the titration of the Spg1p GTPase off the SPB (Li et al. 2000). This suggests that the cellular levels of both Bfa1 and Byr4p could similarly be important to ensure timely mitotic exit or cytokinesis respectively. Interestingly, in *S. pombe* the cellular levels of Byr4p and Spg1p appear to be tightly coordinated, and proteasomal degradation of Byr4p is initiated if Byr4p and Spg1p are not complexed within the cell (Krapp et al., 2008). Although the stability of Bfa1 does not appear to be regulated across the cell cycle, the phosphorylation state of the protein changes dramatically with cell cycle stage (Hu et al., 2001). Specifically, phosphorylation by both Kin4 and Cdc5 are important to activate or inhibit the GAP activity of Bfa1 respectively (Hu et al., 2001; Geymonat et al., 2003; Maekawa et al., 2007). Thus in the budding yeast system, phosphorylation status, rather than proteolysis, regulates Bfa1-Bub2 GAP activity. This allows the GAP activity of Bfa1 to be regulated independently of its positive role in mediating Tem1 localization.

### Additional positive role for Bfa1 in cytokinesis regulation

Our findings indicate that *tem1Δ CDC15-UP* cells have significant cytokinesis defects. These defects appear to be caused by the absence of *TEM1*, rather than the hyperactivation of *CDC15*, since cells containing *CDC15-UP* alone do not exhibit chained cells (Figure 7B-D). Tem1 has a poorly understood role in cytokinesis that is independent of its role in promoting mitotic exit in anaphase. During cytokinesis, the septin ring splits, which allows for actomyosin ring (AMR) contraction, primary and secondary septum formation, and septum degradation in the daughter cell. This process leads to abscission between the mother and daughter cells (reviewed in Juanes and Piatti, 2016). Septin ring splitting occurs before AMR contraction during cytokinesis (Lippincott et al. 2001). It was previously found that in *GAL-UPL-TEM1 net1-1* cells, where the mitotic exit requirement for Tem1 has been bypassed by mutation of the Cdc14 inhibitor *CFI1/NET1*, both septin ring splitting and AMR contraction are defective under TEM1-depletion conditions (Lippincott et al., 2001). Another cytokinesis activator Cyk1/Iqg1 (IQGAP) is dispensable for septin ring splitting, but is required for activation of AMR contraction. Specifically, the GAP-related domain (GRD) of Cyk1/Iqg1 is required for the protein to activate AMR contraction. Interestingly, Tem1 binds specifically to the GRD of the cytokinesis activator Cyk1/Iqg and this raises the possibility that Tem1 regulates this protein’s activity (Shannon and Li, 1999). How this regulation is achieved and where in the cell it occurs is not understood, since Cyk1/Iqg1 localizes to the AMR at the budneck while Tem1 is not visible at this structure.

We found that the cytokinetic defects observed in *tem1Δ CDC15-UP* cells are suppressed by the overexpression of *BFA1.* Specifically, both the Dbf2 budneck localization defects, as well as the chained cell morphology defects, were ameliorated by *GAL-BFA1* (Figure 7). Therefore, our work shows for the first time that *BFA1* activates the process of cytokinesis independently of *TEM1.* Most likely, the positive role for Bfa1 in cytokinesis is through Mob1-Dbf2. Mob1-Dbf2 localization to the budneck is a requirement for actomyosin ring (AMR) contraction and for primary septum formation. Specifically, the F-BAR protein Hof1 is a direct target of Mob1-Dbf2 and thereby promotes AMR constriction (Frenz et al., 2000; Meitinger et al., 2010; Meitinger et al., 2011; Oh et al., 2012; Meitinger et al., 2013). In addition, we found that *tem1Δ CDC15-UP* cells containing a Mob1-eGFP fusion do not exhibit defects in Mob1-eGFP localization to the budneck, nor do they display cytokinesis defects (data not shown). However, the *MOB1-eGFP* allele exhibits some hyperactivity, as evidenced by the localization of this protein to SPBs in metaphase (AS, unpublished observations). Therefore, we conclude that despite having normal Dbf2 kinase activation and cell cycle kinetics, *tem1Δ CDC15-UP* cells have defects in directing the active Mob1-Dbf2 complex to the budneck to fulfill its cytokinetic role (Rock and Amon, 2011; Figure 7). The accurate targeting of Mob1-Dbf2 during cytokinesis is enhanced by *GAL-BFA1* (Figure 7A).

What could be the role of Bfa1 in promoting proper cytokinesis? In *tem1Δ CDC15-UP* cells, the polo kinase Cdc5 is required for mitotic exit (Rock and Amon, 2011). Recently it was shown that Bfa1 is required for the outer plaque dSPB localization of Cdc5 (Botchkarev et al., 2017). The role of Cdc5 localization to dSPBs is likely three-fold: 1) the phosphorylation and inhibition of Bfa1 GAP activity (Hu et al., 2001; Geymonat et al., 2003); 2) the phosphorylation and activation of the essential Cdc15 kinase (Rock and Amon, 2011); and 3) the initiation of septum formation during cytokinesis. To highlight the evidence in support of the role of Cdc5 dSPB localization in the activation of septum formation, previous studies showed that the *CDC5ΔC-CDC12* allele produced a mutant fusion protein that localized to the budneck, but not to SPBs (Park et al., 2004). This mutant allele was capable of initiating mitotic exit in cells lacking *BFA1.* However, the Cdc5ΔC-Cdc12 mutant protein was defective in initiating septum formation and cytokinesis, exhibiting chained cell growth. In contrast, the *CDC5ΔC-CNM67* allele, which localized to SPBs but not to the budneck, exhibited proper mitotic exit, as well as efficient septum formation and cytokinesis. The authors concluded that Cdc5 SPB localization was required for efficient cytokinesis (Park et al., 2004). More recently, it was shown that Cdc5 inhibits the GTPase Cdc42 in late anaphase, which was also required for efficient septum formation (Atkins et al., 2013). However, it is not known whether the outer plaque SPB localization of Cdc5 is necessary for inhibition of Cdc42. In addition, it is not yet known whether the effects of Cdc5 on cytokinesis occur in a Mob1-Dbf2 dependent manner. In this context, it would be important to examine whether *GAL-BFA1* cells display either decreased or increased recruitment of Cdc5 to the dSPB in order to determine whether the effects of *GAL-BFA1* on cytokinesis are mediated through Cdc5. While our results show that Dbf2 recruitment to the primary septum is reduced in the absence of *TEM1*, the links between Bfa1, Cdc5, and Mob1-Dbf2 function in cytokinesis have not been explored.

What is the significance of this dual positive and negative role for Bfa1 on Cdc14 regulation and mitotic exit? Why does Bfa1 have an additional positive function in cytokinesis regulation? One possible explanation is that the use of these same cellular components for multiple cell cycle functions allows for the temporal coupling of mitotic exit and cytokinesis (reviewed in Seshan and Amon, 2004). This is critical in order for cells to restrain cytokinesis until the completion of chromosome segregation. In the *S. pombe* system, the SIN pathway regulates septation and cytokinesis, but not mitotic exit (reviewed in Krapp and Simanis, 2008). Consequently, the deletion of Byr4p, the Bfa1 homolog in fission yeast, causes lethality due to resulting hyperactivity of Spg1p followed by multiple rounds of septation. Thus, in the absence of *byr4*, the process of septation is completely uncoordinated with other mitotic events (Song et al., 1996). Importantly, this is because cells lacking Byr4p are still capable of localizing Spg1p to SPBs, which allows for hyperactivation of the SIN (Sohrmann et al., 1998; Cerutti and Simanis, 1999). In the absence of *BFA1* in *S. cerevisiae*, Tem1 localizes to both SPBs later in anaphase than in wild type cells, and cell cycle progression is normal in an unperturbed cell cycle (Pereira et al., 2000; Valerio-Santiago and Monje-Casas, 2011). Perhaps this is due to the fact that, as we have shown here, Bfa1 is a key anchor for Tem1 at SPBs. The weaker localization of Tem1 in the absence of *BFA1* may safeguard budding yeast cells from MEN hyperactivity and prevent *bfa1Δ* lethality. Further investigation of the role of Bfa1 as a scaffold at the dSPB will lead to clarification of the role of this multi-functional protein in the efficient and accurate execution of late M phase events.

### MATERIALS AND METHODS

#### Yeast strains and growth conditions

All yeast strains used in this study were derivatives of W303 (AS3). The *CDC15-UP* construct was described in Rock and Amon (2011). Culturing conditions were described in the figure legends.

#### Fixed-cell imaging

Indirect immunofluorescence was performed as described previously for α-tubulin (Tub1) and α-HA to detect Cdc14-HA (Visintin *et al.*, 1999). Images for Figures 1B, 2, 4A-F, 5, and 6 were acquired using a Zeiss Axioplan 2 (Zeiss, Thornwood, NY) with a Hamamatsu Orca-R2 camera (Hamamatsu, Middlesex, NJ) and a 63x objective. Cells for Figures 3 and 4G were fixed in a 4% paraformaldehyde, 3.4% sucrose solution for 15 minutes. Cells were washed once in potassium phosphate/sorbitol buffer (1.2M sorbitol, 0.1M potassium phosphate, pH 7.5), and treated with 1% Triton X-100 for 5 minutes. Cells were washed again in potassium phosphate/sorbitol buffer and resuspended in potassium phosphate/sorbitol buffer containing 4’, 6-diamidino-2-phenylindole dihydrochloride before imaging. Cells for Figure 7B-C were fixed in 3.7% formaldehyde, 0.1M potassium phosphate, pH 6.4 prior to sonication and imaging. These cells were imaged using a DeltaVision Elite microscope (GE Healthcare Bio-Sciences, Pittsburgh, PA) with an InsightSSI solid-state light source, a CoolSNAP HQ2 camera, and a 60x plan-ApoN objective.

#### Live-cell imaging

Cells for Figure 7A were imaged directly from log phase cultures using a DeltaVision Elite microscope (GE Healthcare Bio-Sciences) with an InsightSSI solid-state light source, a CoolSNAP HQ2 camera, and a 60x plan-ApoN objective.

## Data availability

All strains are available upon request. The authors affirm that all of the data necessary for confirming the conclusions made within this article are contained within the article and its figures.

## ACKNOWLEDGEMENTS

We thank Angelika Amon and Ian Campbell for providing a critical reading of the manuscript. We are grateful to Angelika Amon for generous sharing of strains and equipment. We also thank Chloe Egan, Aydah Mwangi and Micah Tilove for technical support.

